# A role for keratins in supporting mitochondrial organization and function in skin keratinocytes

**DOI:** 10.1101/822403

**Authors:** Kaylee Steen, Desu Chen, Fengrong Wang, Song Chen, Surinder Kumar, David B. Lombard, Roberto Weigert, Abigail G. Zieman, Carole A. Parent, Pierre A. Coulombe

## Abstract

Mitochondria fulfill essential roles in ATP production, metabolic regulation, calcium signaling, generation of reactive oxygen species (ROS) and additional determinants of cellular health. Recent studies have highlighted a role for mitochondria during cell differentiation, including in skin epidermis. The observation of oxidative stress in keratinocytes from *Krt16* null mouse skin, a model for pachyonychia congenita (PC)-associated palmoplantar keratoderma, prompted us to examine the role of Keratin (K) 16 protein and its partner K6 in regulating the structure and function of mitochondria. Electron microscopy revealed major anomalies in mitochondrial ultrastructure in late stage, E18.5, *Krt6a/Krt6b* null embryonic mouse skin. Follow-up studies utilizing biochemical, metabolic, and live imaging readouts showed that, relative to controls, skin keratinocytes null for *Krt6a/Krt6b* or *Krt16* exhibit elevated ROS, reduced mitochondrial respiration, intracellular distribution differences and altered movement of mitochondria within the cell. These findings highlight a novel role for K6 and K16 in regulating mitochondrial morphology, dynamics and function and shed new light on the causes of oxidative stress observed in PC and related keratin-based skin disorders.

## Introduction

Keratinocytes, the primary cell type that constitutes the skin epidermis, must be able to proliferate, move unidirectionally, assemble and remodel strong adhesive sites as they differentiate and resist mechanical stress. A major family of proteins that impacts all of these functions are the keratin (K) intermediate filaments (IFs), which are regulated in a tissue type-, differentiation-dependent and context-specific manner [1,2]. The human genome features 54 distinct and conserved keratin genes that are partitioned into type I and type II subtypes based on IF sequences [3]. Type I and II keratin genes are expressed in a pairwise fashion and their protein products interact obligatorily to form heteropolymeric 10-nm wide IFs [2]. Mutations in keratin genes have been linked to a variety of skin disorders exhibiting a broad range of tissue level and cellular phenotypes [4]. One such keratin pair, the type II K6 and type I K16, is primarily expressed in ectoderm-derived epithelial appendages in adult skin under normal conditions [2,4,5]. The mature interfollicular epidermis, interestingly, does not express either K6 or K16 unless it experiences injury, UV exposure, or other stresses [4,6,7]. Mutations in either *KRT6A*, *KRT6B*, *KRT6C* or *KRT16* (usually, dominantly-acting missense alleles) can cause PC [8–10]. The most clinically significant aspect of PC is palmoplantar keratoderma (PPK), which is acutely painful and presents as thick calluses developing in the palms and especially soles as a result of oxidative stress and misregulated innate immunity and epidermal homeostasis [6,11].

In addition to forming and functioning as filamentous networks, keratins are known to regulate signaling pathways through protein-protein interactions and modulate organelle processes [12]. For instance, there is an emerging connection between mitochondrial biology and IFs that has the potential to alter the cellular levels of ROS and metabolic flux [12–14]. As mentioned above, our laboratory has previously reported that normal redox balance requires functional K16 via Nrf2 activation and glutathione synthesis. In the case of PC-related PPK, as modeled in *Krt16 null* mice, an imbalance in redox homeostasis precedes the appearance of lesions and can be rescued using a Nrf2 activator [11]. These observations prompted us to investigate mitochondria as a source of dysfunction since this organelle is a major hub of ROS production and is key to cell energetics and redox homeostasis. Besides, we previously reported on ultrastructural anomalies in late embryonic skin of *Krt5* null mice [15] whereas others have described that K8, a type II keratin expressed in simple epithelia, modulate mitochondrial network formation [13,14]. Here, we report that lack of K16 and, to a greater extent, lack of K6 impair mitochondrial cristae formation, respiration, and dynamics in skin keratinocytes. These findings suggest that disruption of mitochondrial-keratin interactions, which in turn leads to impaired cellular and redox homeostasis, is related to oxidative stress that precede PPK lesions.

## Results

As previously reported, mice homozygous for a *Krt6a/Krt6b* double-null allele are born with the expected frequency but rapidly develop oral lesions that hampers their postnatal growth and results in their untimely death within a week post-birth [16]. We used transmission electron microscopy (TEM) to compare the ultrastructural features of epidermis and mitochondria in particular in late stage (E18.5) embryonic *Krt6a/Krt6b* null and *WT* back skin. Low magnification surveys of epoxy-embedded tissue sections show that the epidermis of *Krt6a/Krt6b* homozygous null mice is intact and shows a normal morphology (Supplemental Figure 1A,B). In contrast, examination at higher magnifications reveal that, relative to control, mitochondria in epidermal keratinocytes lacking both keratins 6a and 6b proteins (K6a/K6b) exhibit several anomalies with respect to shape and cristae organization (Figure 1A-D). The mitochondria in mutant skin exhibit a swollen appearance with little to no intact cristae (Figure 1B,D), in contrast to the elongated and electron-dense morphology seen in normal control tissue (Figure 1A,C). To quantify these observations, we categorized mitochondria as intact (cristae running all the way through a single mitochondria), partially abnormal (some cristae observed but not fully intact), and severely abnormal (no cristae) (note: representative images are shown in Supplemental Figures 1C-E). This confirmed that significantly more mitochondria show a severely abnormal ultrastructure compared to control tissue (Figure 1E). While these anomalies could be the result of inefficient fission and/or fusion leading to disorganized cristae (Figure 1B,D) [17–19], we did not observe any major expression differences in relevant biomarkers (*Mfn1*, *Mfn2*, *Opa1* and *Drp1*) with the exception of a small increase in *Mfn2* mRNA in *Krt6a/Krt6b null* keratinocytes compared to *WT* cells cultured in primary conditions (Figure 1F). We observed a modest trend towards lower mRNA levels for the mitochondrial markers cytochrome c oxidase subunit 4 (COX4), succinate dehydrogenase (SDH), translocase of the inner membrane (Tim23), complex III subunit 2 (COR2), and pyruvate dehydrogenase (PDH) in *Krt6a/Krt6b null* keratinocytes relative to *WT*, with Tim23 and PDH reaching statistical significance (Figure 1G). However, there were no differences observed at the protein level with these markers (data not shown), suggesting that fission and fusion processes do not directly contribute to mitochondrial defects. Finally, we did not observe any change in mitophagy markers responsible for mitochondrial turnover (data not shown), which is consistent with data gathered from keratinocytes of PC patients with K6a mutations ([10]; see Discussion). Of note, we previously reported on the occurrence of similar ultrastructural defects in mitochondria from *Krt5 null* E18.5 mouse epidermis [15]. The loss of another IF protein, desmin, was also found to induce mitochondrial swelling and matrix disruption in cardiac muscle. Importantly, the mitochondria in desmin-null cardiac muscle display a very similar phenotype to that seen in *Krt6a/Krt6b* null skin (see Figure 1B and [20]). This suggests that, as is the case for other IFs, K6- and/or K16-containing filaments likely plays a direct role in mitochondrial architecture.

**Figure 1.**
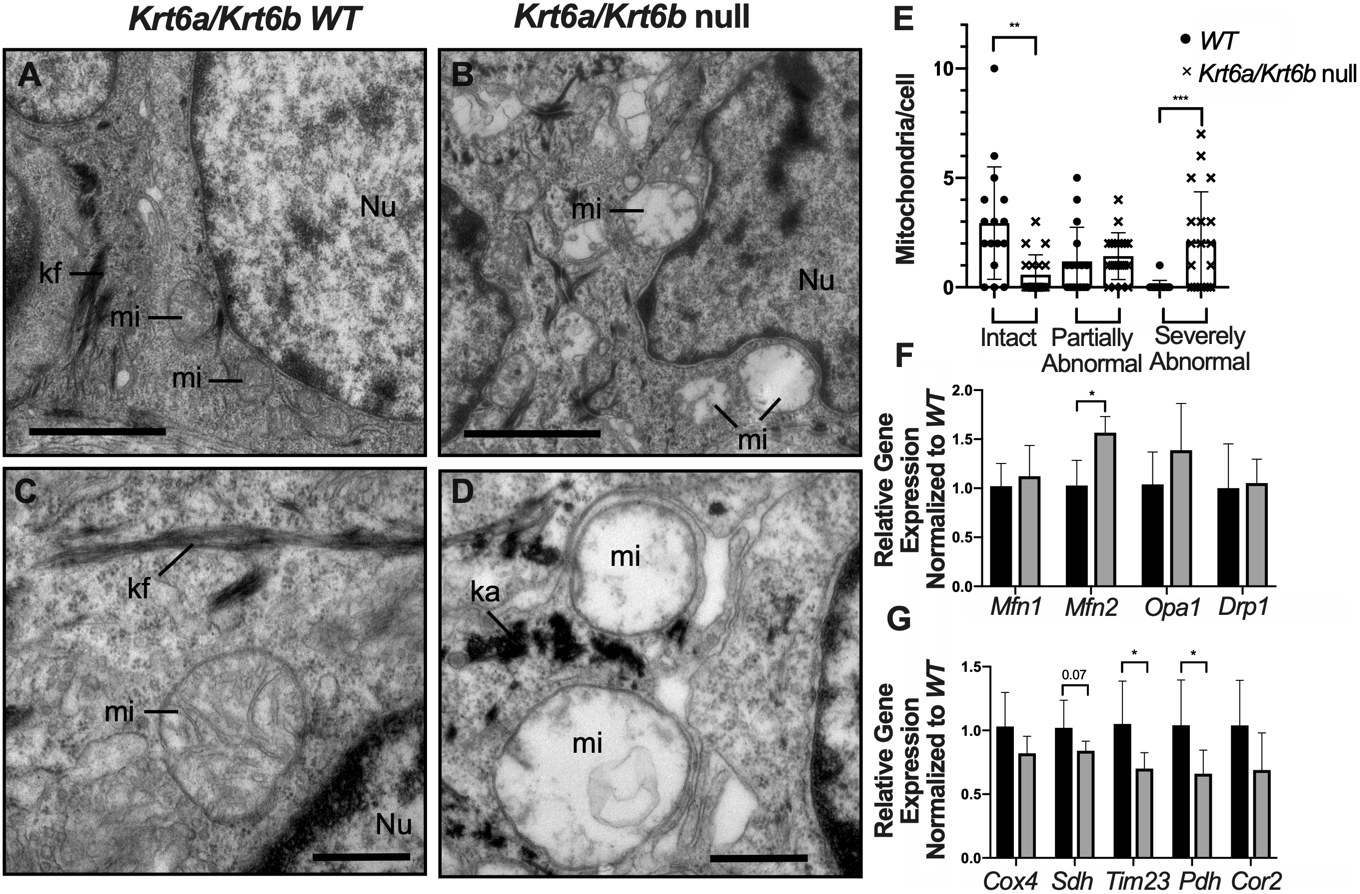
Transmission electron microscopy of E18.5 back skin. (A-D) Six pups were analyzed from *WT* and *Krt6a/Krt6b* null littermates. Images are shown at lower (A-B; bar equals 2 μm) and higher magnification (C-D; bar equals 500 nm) for each genotype. ka, keratin aggregates; kf, keratin filaments; mi, mitochondrion; Nu, nucleus. (E) Quantification of the frequency of mitochondria per cell that appeared intact, partially abnormal or severely abnormal (representative images of each category are shown in Supplemental Figure 1 C-E). (F-G) Analysis of specific mRNA transcripts in total RNA samples. Keratinocytes were isolated from P1 *WT* and *Krt6a/Krt6b* null littermates and seeded at equal cell density for culture. Total RNA was isolated and analyzed. Gene expression of mitochondrial markers was normalized to 18S RNA in each sample. n=6 and 3 independent experiments were performed. Student’s t test was used with significance set at p<0.05.

Keratin proteins physically interact with several cellular organelles [13,14,21,22], including mitochondria. We next compared the subcellular distribution of K16, for which we have a high titer (monospecific) antibody [5,23], to that of PDH using high resolution confocal microscopy (Airyscan technology) in *WT* newborn mouse skin keratinocytes in primary culture. The signal for K16 closely aligns with that of PDH, which otherwise is polarized towards the perinuclear space (Figures 2A-C). When using conventional confocal microscopy, we find that the absolute spatial distribution of the mitochondria (as measured by PDH) is altered when either K6 or K16 are missing. Relative to the perinuclear localization in *WT* control keratinocytes in primary culture, the PDH signal is broadly distributed throughout the cytoplasm in *Krt6a/Krt6b* null keratinocytes, with a higher PDH signal at the cell periphery (Figure 2D,E; quantitation in Figure 2F). This effect was also observed in spontaneously immortalized keratinocytes (SIMEKs) lacking *Krt16* compared to *WT* SIMEKs (data not shown). These findings suggest that K6- and/or K16 – containing filaments may directly or indirectly impact the organization of mitochondria in skin keratinocytes. Loss of K6 or K16 leads to a differential dispersion of mitochondria throughout the cell, with the potential to disrupt the physiological signaling capacity of this organelle.

**Figure 2.**
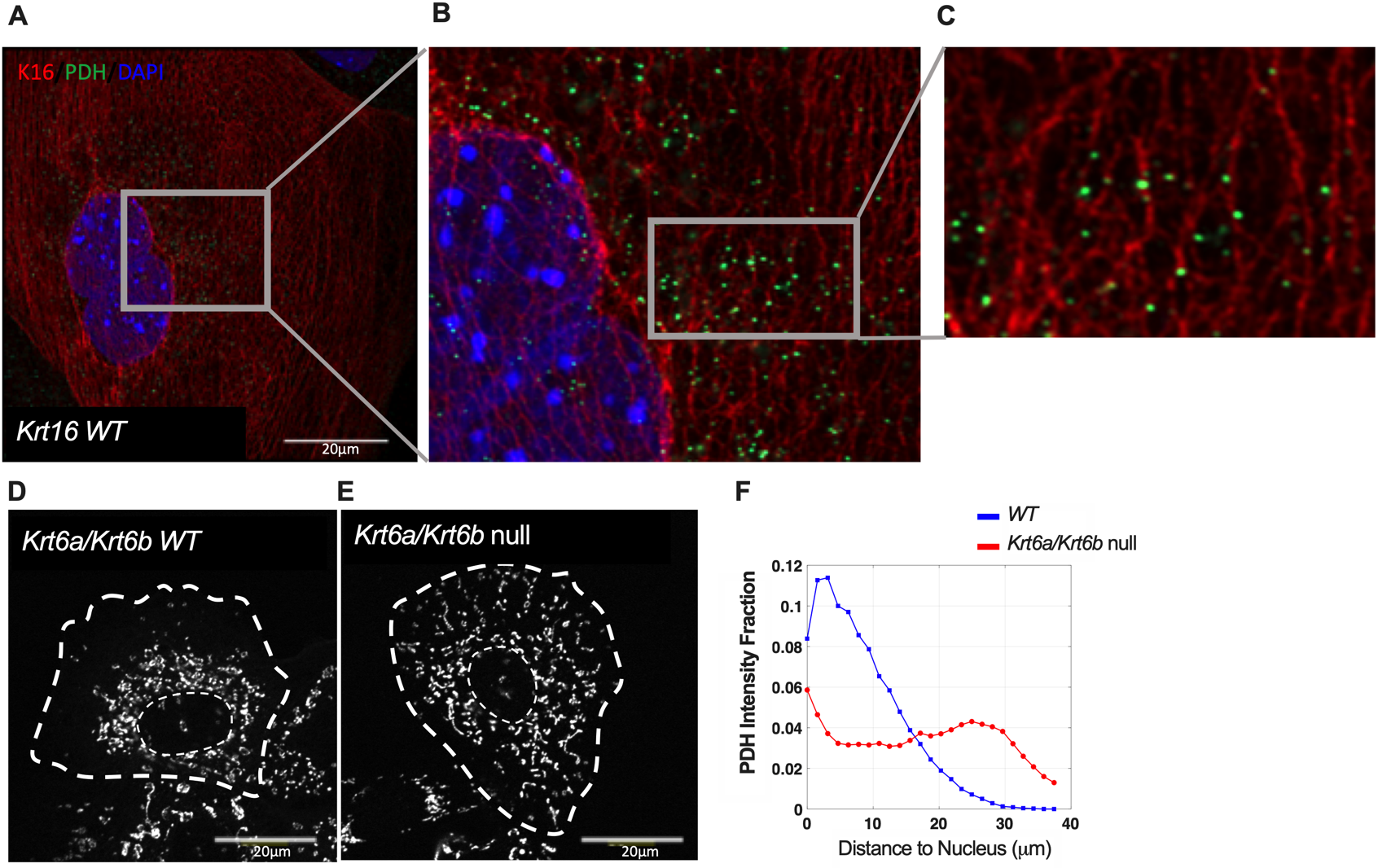
Indirect immunofluorescence for K16 and pyruvate dehydrogenase (PDH) in keratinocytes in primary culture. (A-C) Keratinocytes were isolated from P1 *WT Krt16* mouse strain and cultured in primary conditions. Images were acquired using an LSM 800 Airyscan mode. Bars equal 20μm. (D-E) Z-stack images of PDH staining in newborn P1 *WT* and *Krt6a/Krt6b null* skin keratinocytes in primary culture. Dashed lines depict the cell periphery and nucleus. (F) Graphs depict PDH signal intensity distribution in the cytoplasm, as the intensity fraction of the total intensity with respect to the distance from nucleus or the distance to the nucleus normalized to the spacing between nucleus and the cell boundary. The curves are the average intensity over 18 *Krt6a/Krt6b null* cells and 34 *WT* cells. Images were acquired using z-stack and maximal projection with an LSM 800 confocal mode. Bars equal 20μm.

Cells rely on the electron transport chain (ETC) of the mitochondria to produce ATP through the reduction and oxidation of ETC protein complexes from electrons donated from the TCA cycle [17]. Efficient operation of this complex process requires that cristae be densely packed inside mitochondria to keep the ETC complexes close together for electron transport [24,25]. If such a continuity is not maintained, electrons can move back into the mitochondrial matrix, react with molecular oxygen and produce excess ROS [17,26,27]. The presence of abnormal mitochondrial ultrastructure led us to hypothesize that the absence of either K6a/K6b or K16 protein leads to imbalances in ROS levels. To test this, we measured total ROS in *WT* and *Krt6a/Krt6b* null keratinocytes cultured under normal and stressed conditions through exposure to tert-butyl hydrogen peroxide (TBHP). Cells lacking K6a/K6b consistently displayed a trend of higher ROS levels relative to *WT* under baseline conditions and this reached statistical significance when stressed with TBHP (Supplemental Figure 2A). A similar trend was observed in *Krt16* null keratinocytes in primary culture, though it did not reach statistical significance (Supplemental Figure 2B). We repeated this assay in spontaneously immortalized cells isolated from *WT* and *Krt16 null* mice and found that, similar to *Krt6a/Krt6b null* ones, immortalized cells lacking K16 displayed higher levels of ROS compared to control cells (Supplemental Figure 2C).

ROS have been implicated in many forms of cellular dysfunction and can directly damage mitochondria [28–30]. To explore how increased ROS affects mitochondrial function in cells lacking either K6a/K6b or K16, we measured mitochondrial respiration using the Seahorse Mito Stress Test Kit (Figure 3A). Keratinocytes isolated from both *Krt6a/Krt6b null* and *Krt16 null* mice (P1) showed a significant reduction in basal and maximal respiration compared to *WT* cells (Figure 3B and 3D). Null keratinocytes also displayed a reduced proton leak (Figure 3C), suggesting that there may be an overall reduction in uncoupling protein activity or increased permeability of the inner mitochondrial, reducing overall electron movement across the ETC [25,31]. Interestingly, maximal respiration for all cell populations did not exceed basal respiration following FCCP treatment. We repeated these experiments, varying the media and drug conditions, but always observed this same results. While this response is not common, it has been previously observed by others [32–34]. This reality suggests that newborn mouse skin keratinocytes in primary cell culture are already maximally respiring even at a resting state. This finding was further supported by reduced membrane potential as measured by TMRE fluorescence in both *Krt6a/Krt6b null* and *Krt16 null* keratinocytes (Figure 3E). Interestingly, other IFs, namely vimentin, have also been shown to maintain membrane potential and loss of this interaction alters mitochondrial positioning and physiological activity [35].

**Figure 3.**
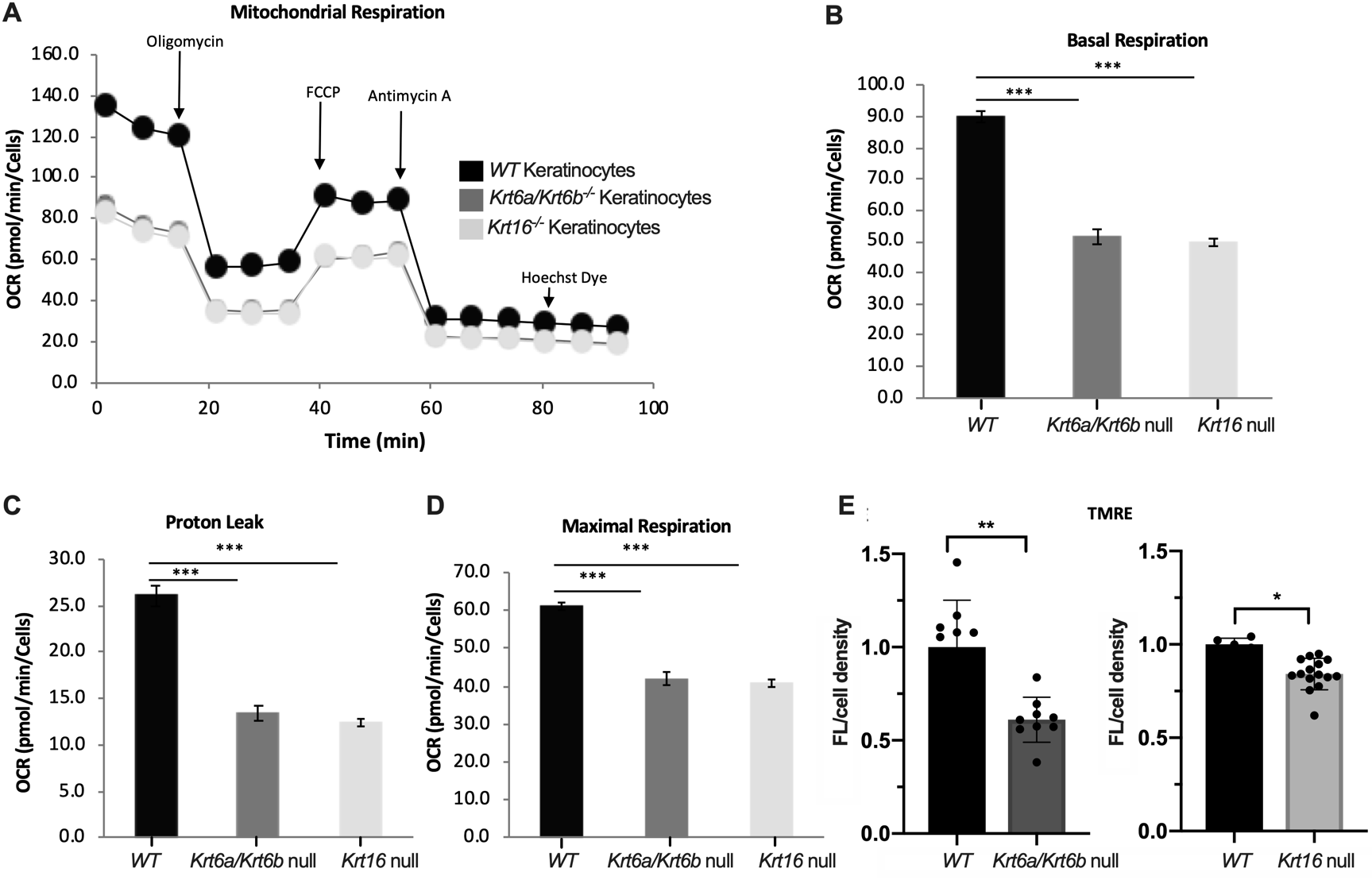
Seahorse analysis of mitochondrial function. 4×10^4^ skin keratinocytes from P1 *WT*, *Krt6a/Krt6b* null and *Krt16* null mice were seeded in primary culture. Analyses (XFe96 instrument) were done using the Agilent protocol for mitochondrial stress test assay (cf. Methods). Samples were normalized by cell density and mitochondrial basal respiration, maximal respiration and proton leak were determined. (A) Mitochondrial respiration was assessed under baseline conditions to measure basal respiration. Oligomycin treatment (ATP synthase inhibitor) was applied to measure proton leak and mitochondrial-linked respiration, FCCP (Carbonyl cyanide-4-phenylhydrazone) treatment (proton uncoupler) to measure maximal respiration and Rotenone/Antimycin A (complex I and III inhibitor) to measure spare respiratory capacity. The *Krt6a/Krt6b* null and *Krt16* null cells each exhibited lower levels of (B) basal respiration, (C) proton leak and (D) maximal respiration with no change in ATP production or spare respiratory capacity (data not shown). n= 10-16 from 3 independent experiments. Student’s t test was used with significance set at p<0.05. (E) Skin keratinocytes were isolated from P1 *WT* and *Krt6a/Krt6b* null littermates (left) or *WT* and *Krt16* null littermates (right) and cultured in primary conditions. 7×10^4^ cells were seeded in a 96-well plate and labeled with 500nM TMRE (Tetramethylrhodamine ethyl ester perchlorate) for 30 minutes, and fluorescence was then measured and normalized to the *WT* controls. Cells were normalized by measuring total DNA using CyQuant dye. n=5-20 from 3 independent experiments performed. Student’s t test was used with significance set at p<0.05.

Mitochondria are dynamic organelles that constantly undergo fission and fusion to achieve an organization that regulates and optimizes several cellular functions in response to exogenous signals [18,36]. Respiration and ROS production are regulated by and can impact mitochondrial dynamics [17,28,37–39]. To assess whether and how K6/K16 alter mitochondria dynamics, we labeled skin keratinocytes in culture (P1 *WT* and null littermates) with MitoTracker Red and performed live cell imaging in real time to monitor mitochondrial movement under normal cellular conditions. In keratinocytes null for either K6a/K6b or K16, mitochondria showed no alteration in size or circularity but exhibited an increased speed of movement [40] (data not shown and Figure 4). Representative findings can be visualized in Supplemental Figure 3, Movies 1-4. A quantitative assessment of the movement of individual mitochondrial profiles, as an overall average (Figure 4A,B) and as a function of mitochondrial size (Figure 4C,D) is reported for *Krt6a/Krt6b* null and *Krt16* null newborn keratinocytes relative to their respective *WT* controls. The increased rate of movement (speed) was unrelated to the size of mitochondrial profiles (represented by the x-axis), although smaller mitochondrial shapes showed a greater rate of movement in the two null genotypes compared to controls (Figure 4C,D). Further, our data conveys a heterogeneity of mitochondrial populations even within the same cell type and individual cells (Supplemental Figure 4). Mitochondria in *Krt6a/Krt6b* null and *Krt16* null keratinocytes also appeared to move in a disorganized or random fashion, but we were unable to quantify this behavior. Overall, these data demonstrate a difference in the movement of mitochondrial profiles in keratinocytes lacking K6a/K6b or K16. While the live imaging performed does not directly measure mitochondrial fission and fusion rates, it extends the findings of disrupted mitochondrial ultrastructure (Figure 1) and reduced respiration (Figure 3), pointing to a state of mitochondrial instability.

**Figure 4.**
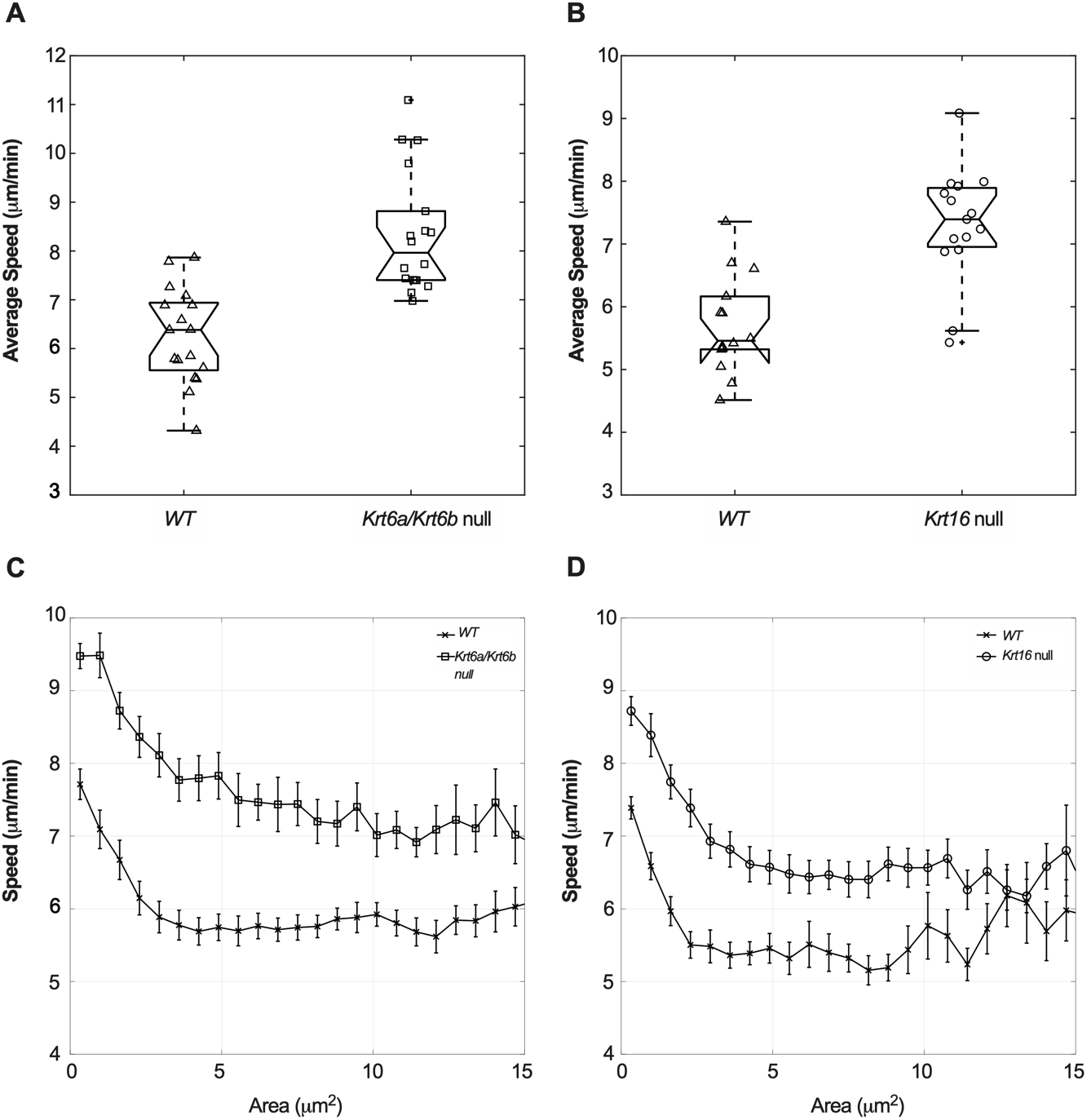
Live imaging of mitochrondrial dynamics. 50,000 skin keratinocytes were obtained from P1 *WT*, *Krt6a/Krt6b null* and *Krt16 null* mice and seeded for primary culture overnight. Cells were then labeled with 50nM of MitoTracker Red and live cell imaging was performed for 5 minutes to track labeled mitochondria of individual cells. Representative movies are shown in Supplemental Videos 1-4. (A-B) Average speed of the tracked mitochondrial shapes with areas between 1 um^2^ and 10 um^2^ in *WT* and *Krt6a/Krt6b null* keratinocytes (A) and *WT* and *Krt16 null* keratinocytes (B). In the box plot, the scatter points represent each cell. The whiskers represent the range of the data and the box indicate the 25% to 75% percentile. The notch with the middle line indicates the median and the cross represents the outlier.). In both box plots, p-value < 0.0001. (C-D) The average speed is tracked as a function of mitochondria size in the *WT* and *Krt6a/Krt6b null* keratinocytes (C) and *WT* and *Krt16 null* keratinocytes (D). There are 3 independent experiments (n > 5 cells for each).

## Discussion

Keratin proteins play a multifaceted role in keratinocyte homeostasis, and mutations in keratin genes lead to a diverse array of phenotypic outcomes. Keratins 6a/6b and 16 are robustly wound-inducible and support and promote a number of cellular functions including structural integrity [16,23,41], cell migration [7,42], keratinocyte differentiation [43], regulation of innate immunity [6] and redox homeostasis [11,44]. Disruption of many of these cellular roles is poised to play a role in the pathophysiology of PC, in particular, oral and palmoplantar keratoderma lesions [45]. Mitochondria represent the main cellular protagonist for regulation of ROS, which it achieves mainly via promoting the integrity and efficiency of the electron transport chain. Silvander *et al*. [13] reported that loss of keratin 8 reduced mitochondrial membrane potential and ATP production in pancreatic b-cells, a novel role that involves an interaction with TCHP. Nishizawa *et al*. [12] provided evidence that K6a/K6b and especially K16 physically interact with TCHP. Here we report that K6a/K6b and K16 each regulate the organization and function of mitochondria in skin keratinocytes. The occurrence of markedly reduced levels of K16 protein in *Krt6a/Krt6b* double-null mouse skin keratinocytes, in both tissue and cell culture settings [7, 16, 41], may help explain the more severe phenotype they exhibit relative to *Krt16* nulls. The novel findings reported here have potential significance for keratinocyte differentiation [38], epithelial homeostasis [10,46,47], and also for the pathophysiology of keratin mutation-based skin epithelial disorders [4,9–11,41]. We note that we could not generate reliable evidence for a physical TCHP-K16 interaction in skin keratinocytes owing in part to the unavailability of a good antibody to TCHP (data not shown).

ROS levels are significantly higher in keratinocytes null for *Krt6a/Krt6b* or *Krt16* relative to *WT*. These findings correlate with reductions in both basal and maximal mitochondrial respiration in the *Krt6a/Krt6b* and *Krt16* ablated states, as measured by Seahorse analysis. The latter also indicated that there is reduced proton leak in the two keratin null settings, which is further supported by the reduced membrane potential prevailing in mitochondria. These findings are consistent with the damaged and disorganized cristae observed in mitochondria of *Krt6a/Krt6b null* epithelial back skin. Besides, mitochondria tend to be localized to the perinuclear cytoplasm in wildtype keratinocytes but show a broader dispersion in keratinocytes lacking K6a/K6b or K16. Of note, keratin filaments themselves readily concentrate to the perinuclear region, particularly in suprabasal keratinocytes of surface epithelia [21,22]. Accordingly, alterations in keratin filament properties resulting from the loss of either K6a/K6b or K16 may prevent the mitochondria from concentrating near the nucleus. The subcellular localization of mitochondria has a major impact on cell signaling and function, including migration, calcium signaling, and gene expression [48–52]. Alteration of this steady state is an exciting area for future studies to determine the associated effects on keratinocyte function. ROS production and mitochondrial function are also closely linked to the latter’s dynamics and network formation, and indeed differences were measured in mitochondria motility in *Krt6a/Krt6b* null and *Krt16* null keratinocytes relative to *WT*. These data suggest that keratins help stabilize mitochondria spatially and structurally in keratinocytes, and that disrupting these processes has significant consequences for cell signaling, bioenergetics, and redox homeostasis [17–19,53].

Our findings significantly extend a recent study showing that mitophagy turnover is impaired in cultures of immortalized keratinocytes derived from individuals with PC [10], thus adding to the evidence that anomalies in mitochondria and in redox balance may play a significant role in the pathophysiology of PC-associated PPK. They also add to a growing body of evidence linking keratin [13] and other types of IFs, notably vimentin and desmin, to mitochondrial regulation and function [20,35,54,55]. Like microtubules and actin filaments, IFs participate in controlling mitochondrial motility, likely through stabilization, which in turn regulates ATP production, calcium signaling and intermediary metabolism [55–58]. Many issues remain unresolved, however. First, while others provided evidence that K16 and K6 can bind to trichoplein, a candidate mitochondrial linker protein [12,13], there is still a need to definitively identify the mechanism in which keratin filaments interact with the mitochondria. Secondly, we do not know whether the mitochondrial dysfunction is leading to increased ROS production or if the damage is a result of an already established oxidative stress. This is significant because activation of oxidative stress pathways precedes PPK lesions in mice [11,44]. The findings reported here provide a platform to answer these questions along with a deeper understanding of keratins’ active role in regulating mitochondrial function, structure and dynamics.

## Materials and Methods

### Mouse handling

All experiments involving mice were reviewed and approved by the Unit for Laboratory Animal Medicine (ULAM) at the University of Michigan. The *Krt6a/Krt6b* null [16] and *Krt16* null [23] mouse strains (C57BL/6 background) were maintained under specific pathogen–free conditions, fed chow and water ad libitum, and bred and genotyped as described. All studies involving E18.5 back skin tissue and newborn skin (P1-P2 pups) keratinocytes in primary culture were performed using littermates with a *WT* or *homozygous null* genotype.

### Reagents

Primary antibodies used include anti-PDH (Abcam), anti-FLAG^®^ M2 Magnetic Beads (Sigma-Aldrich), anti-TCHP (abcam), anti-b-actin, anti-K16 [5] and anti-K6 [4]. MitoTracker CMXRos (ThermoFisher) was used for live cell imaging of mitochondria. Secondary antibodies used included HRP-conjugated secondary antibodies (Sigma-Aldrich) and Alexa Fluor 488, Alexa Fluor 594 and Alexa Fluor 647 (ThermoFisher).

### Tissue preparation for sectioning and microscopy

*Krt6a/Krt6b* heterozygous mating pairs were set up and E18.5 pups were harvested and genotyped [59]. Tissue sectioning for immunostaining was performed by submerging back skin into OCT (Sakura Finetek), freezing at −20°C, and preparing 5uM sections using CryoStar NX50 (Thermo Scientific), and stained as described below. For transmission electron microscopy, E18.5 back skin from *WT* and *Krt6a/Krt6b null* littermates were placed flat at the bottom of a 24-well plate and treated with 1mL of fixative (2.5% glutaraldehyde, 3% paraformaldehyde in Sorenson’s buffer). The samples were then given to the University of Michigan’s Microscopy and Image Analysis Laboratory (MIL) core for thin-sectioning. Sections were imaged using the JEOL 1400 Plus TEM at the MIL.

### Keratinocyte culture

*WT* and *Krt6a/Krt6b null* and *WT* and *Krt16 null* littermates were taken at P1. The skin was removed and left in 0.25% Trypsin overnight at 4°C, and following day keratinocyte isolation was performed as described [7]. Cells were counted and seeded on collagen I coated plates in differentiation-promoting mKER media for two days unless specified otherwise. Skin-derived keratinocytes express K6 and K16 protein under these circumstances [4,5,7,60]. Spontaneously immortalized mouse epidermal keratinocyte (SIMEK) cell lines were generated from *Krt16* null and *WT* littermates as described in Reichelt *et al*. [60].

### Mitochondrial respiration

Mitochondrial stress test was performed to measure oxygen consumption rate (OCR), using the XFe96 Extracellular Flux Analyzer (Seahorse Bioscience, Billerica, MA, now Agilent Technologies, Santa Clara, CA), as per manufacturer’s instructions. Briefly, 4×10^4^ keratinocytes from the back skin of *WT* and *Krt6a/Krt6b* null, and *WT* and *Krt16* null, littermates were plated in complete mKER media supplemented with 10% (v/v) heat-inactivated FBS into each well of 96-well Seahorse microplates. Cells were then incubated in 5% CO_2_ at 37°C for 24 hours. Following incubation, cells were washed twice, incubated (in non-CO_2_ incubator at 37°C), and analyzed in XF assay media (non-buffered DMEM containing 25 mM glucose, 2 mM L-glutamine, and 1 mM sodium pyruvate, pH 7.4) at 37°C, under basal conditions and in response to 1 μM oligomycin, 2 μM fluoro-carbonyl cyanide phenylhydrazone (FCCP) and 0.5 μM rotenone + 0.5 μM antimycin from the Seahorse XF Cell Mito Stress Test Kit (Agilent Technologies). Data was analyzed by the Seahorse XF Cell Mito Stress Test Report Generator. OCR (pmol O_2_/min) values were normalized to the cell count using Hoechst dye in the final prot injection and transferred to the Cytation 5 (BioTek, Winooski, VT).

### Measurement of reactive oxygen species

Keratinocytes were cultured in primary conditions from *WT* and *Krt6a/Krt6b null*, and *WT* and *Krt16 null* littermates. Cells seeded in the Corning™ 96-Well Clear Bottom Black Polystyrene Microplates at a cell density of 5×10^4^. Due to the fast-growing nature of the immortalized cells, 2.5×10^4^ cells were seeded for the *WT* and *Krt16 null* SIMEKs. The protocol from Abcam’s DCFDA cell ROS detection assay kit was used to measure total cellular ROS at baseline and with increasing concentrations of Tert-Butyl Hydrogen Peroxide.

### Measurement of mitochondrial membrane potential

Keratinocytes were isolated from *WT* and *Krt6a/Krt6b null* and *WT* and *Krt16 null* littermates and seeded in the Corning™ 96-Well Clear Bottom Black Polystyrene Microplates at a cell density of 7×10^4^. Following 48 hours of culturing, cells were incubated with 500nM of TMRE dye for 30 minutes followed by two washes with PBS containing 0.2% BSA. Fluorescence was measured at Ex/Em = 549/575 nm and normalized to cell number using the CyQUANT kit (ThermoFisher). Normalization was done by aspirating the media and freezing the cells at −80°C overnight. The cells were then labeled with CyQuant dye to measure total DNA.

### Analysis of mitochondrial dynamics

Live cell imaging was performed to track the mitochondrial dynamics in *WT*, *Krt6a/Krt6b null* and *Krt16 null* keratinocytes. Cells were seeded in Collagen I coated wells at a density of 50K cells to reach about 60-70% confluency in order to image individual cells (Nunc™ Lab-Tek™ II Chambered Coverglass with a No. 1.5 borosilicate glass bottom for 48hrs). Prior to imaging, cells were incubated in Phenol red-free DMEM 0.5% BSA media containing 50nM of MitoTracker red for 15 minutes at 37°C. Cells were then washed once with Phenol red-free DMEM 0.5% BSA media and imaged using the Zeiss Airyscan LSM 880. Videos were taken with a 0.11 laser power, 0.5ms intervals, and 500 cycles with Airyscan processing. Due to imaging limitations, we were not able to resolve individual mitochondria. The objects segmented and measured most likely represent clusters of mitochondria (mitochondrial shapes). For each time frame mitochondrial shapes were segmented using the snake algorithm, an active contour model implemented with custom MATLAB scripts. In brief, this algorithm defined the outlines of mitochondrial shapes by a set of representative points along the boundary, i.e., “snakes” [61]. Snake contours were initiated with the convex hulls of the objects and converged to fit the boundary in the energy minimization iterations defined by force fields determined by brightness gradient of the fluorescence. Parameters used to identify outlines of mitochondrial shapes were: 1) the snake elasticity parameter α (0.01), 2) the viscosity parameter γ (0.05), and 3) the external force weight k (0.5). Two mitochondrial shapes were considered the same object if they overlapped for more than 60% in two successive frames. Mitochondrial shapes that appear to fragment or fuse in successive frames were excluded from the quantification. Speed of the center of mass and the area of the tracked objects were determined.

### Indirect immunofluorescence and quantitation

Immunostaining (indirect immunofluorescence) was performed on keratinocytes and epidermal back skin tissue sections. Samples were fixed in 4% paraformaldehyde for 10 minutes at room temperature, washed in PBS, blocked in 5% normal goat serum/0.1% Triton X-100/ PBS for 1 hour at room temperature, incubated in primary antibody solution overnight at 4°C, washed in PBS, incubated in secondary antibody solution for 1 hour, incubated with DAPI for 5 minutes, washed in PBS, and mounted in Fluor-Save Reagent Mounting Medium (EMD Millipore) before visualization using a Zeiss LSM 800 fluorescence confocal microscope. To quantify the mitochondrial PDH distribution throughout the cell, the nucleus and the cell boundaries were manually segmented with Fiji. Each boundary loop was then smoothed in MATLAB and 100 points on the boundary with even spacing were picked to partition the loop into 100 sections. The points on the cell boundary were linked to corresponding points on the cell nucleus boundary, keeping the circular order of the points and the total distance between paired points at the minimum. In this way, the area between the nucleus and the cell boundary was segmented into 100 regions. For each region (and the associated boundary points) the local distance between the nucleus and the cell boundary was defined as the distance between the middle points of the local nucleus and cell boundary sections. The distance of each pixel in the segmented regions to the nucleus was defined as the distance between the pixel to the middle point of the local nucleus boundary section. As some cells touched other cells, only the regions with cell boundary not touching other cells were manually selected for further analysis. To quantify the average PDH fluorescence intensity with respect to the distance to the nucleus, the distance was binned every 10 pixels (1.6 μm). The average PDH fluorescence intensity of pixels in each distance bin was then calculated.

### Biochemical analyses

RNA was isolated from primary keratinocytes cells using the NucleoSpin RNA kit (Macherey-Nagel) followed by cDNA preparation (iScript cDNA synthesis kit; Bio-Rad). All oligonucleotide primers (see Supplemental Table 1) were designed using the *mus musculus* RefSeq through the NCBI database. K16-flag protein (C-terminally tagged) was made using the pcDNA™3.1 Directional TOPO™ Expression Kit (ThermoFisher) using primers for the mouse *Krt16* gene with flag peptide sequence on the 3’ end (Supplemental Table 1).

### Statistical analyses

All statistical analyses, unless indicated elsewhere, were performed using a two-tailed Student’s *t* test.

## Supporting information

Steen et al. Supplemental Information

Video 1 - Steen et al (2020)

Video 2 - Steen et al (2020)

Video 3 - Steen et al (2020)

Video 4 - Steen et al (2020)

## Acknowledgements

We thank Samarth Setru and Ritankar Majumdar for assistance. These studies were supported by R01 grants AR044232 and AR042047 (to P.A. Coulombe), R01 grant GM101171 (to D.B. Lombard), and T32 grant AR007197 (trainee support for K. Steen) from the National Institutes of Health.

---

**Supplemental Figure 1.** Morphological analyses (related to Figure 1). (A-B) Low-magnification surveys of epoxy-embedded E18.5 back skin tissue sections (1 um thick) from *WT and Krt6a/Krt6b null* mouse littermates. Bars equal 50um. Epi, epidermis; HF, hair follicles. (C-E) Representative images of (C) intact mitochondria (cristae running all the way through the organelle), (D) partially abnormal mitochondria (some cristae observe but contain large gaps or disorganized) and (E) severely abnormal mitochondria (little to no cristae observed). Bars equal 500nm.

**Supplemental Figure 2.** ROS measurements. (A) Skin keratinocytes were isolated from P1 *WT* and *Krt6a/Krt6b* null littermates and 5×10^4^ cells were seeded in a 96-well plate for primary culture. Cells were given fresh media the following day, and labeled 24 hours later with DCFDA dye (2’,7’ – dichlorofluorescin diacetate abbreviated as “D”; 0-25 μm) and Tert-Butyl Hydrogen Peroxide (abbreviated as “T”; 0-250 μm) to induce oxidative stress. ROS were measured using the Abcam DCFDA cellular ROS detection assay kit. (B) Keratinocytes were cultured from *WT* and *Krt16* null littermates and cultured and analyzed as described in (A). (C) Spontaneously immortalized keratinocytes from P1 *WT* and *Krt16* null mice were seeded at a density of 2.5×10^4^ cells per well and assayed as described in (A). n= 3 independent experiments. Student’s t test; significance at p<0.05.

**Supplemental Figure 3** Representative movies of fluorescently-labeled mitochondria in primary cultures of keratinocytes (related to Figure 4). *WT* (Movie 1) and *Krt6a/Krt6b* null (Movie 2) newborn mouse littermates; *WT* (Movie 3) and *Krt16* null (Movie 4) newborn mouse littermates. The quantitative data reported in Figure 4 was obtain from analyzing such movies.

**Supplemental Figure 4.** Complement to live imaging of mitochondrial dynamics (related to Figure 4). Data corresponds to the average mitochondrial speed as a function of mitochondrial profile size for each of three independent experiments involving matched null and WT littermates for each of the *Krt6a/Krt6b* and *Krt16* null lines.

**Supplemental Table 1.**
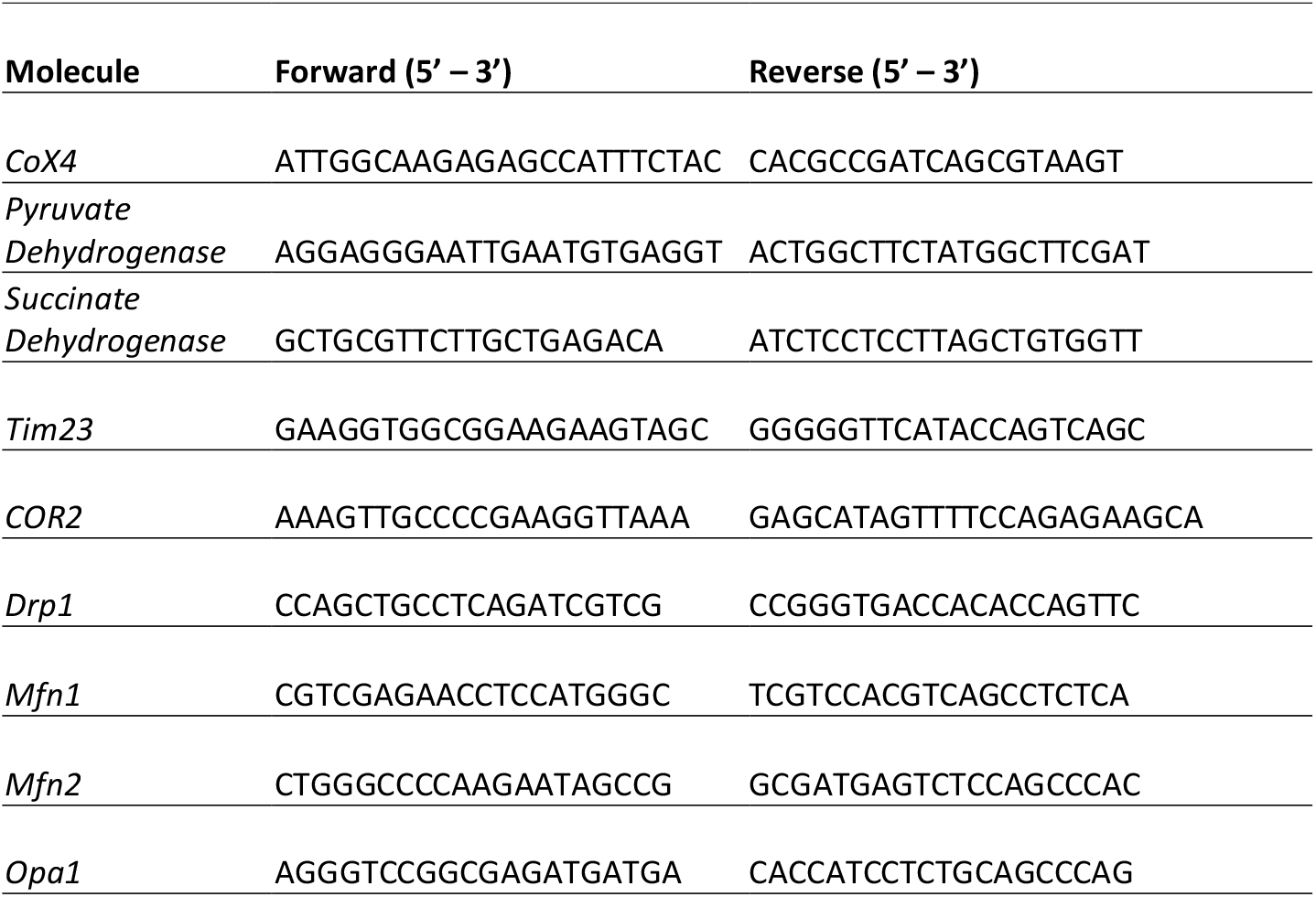
Forward and reverse oligonucleotide primers used for RT-qPCR assays (listed in order they appear in the text).

