## Supplementary material for "A role for keratins in supporting mitochondrial organization and function in skin keratinocytes": Steen et al. Supplemental Information

**List of elements:**

**Supplemental Figure 1.** Morphological analyses (related to Figure 1). (A-B) Low-magnification surveys of epoxy-embedded E18.5 back skin tissue sections (1 um thick) from *WT and Krt6a/Krt6b null* mouse littermates. Bars equal 50um. Epi, epidermis; HF, hair follicles. (C-E) Representative images of (C) intact mitochondria (cristae running all the way through the organelle), (D) partially abnormal mitochondria (some cristae observe but contain large gaps or disorganized) and (E) severely abnormal mitochondria (little to no cristae observed). Bars equal 500nm.

**Supplemental Figure 2.** ROS measurements. (A) Skin keratinocytes were isolated from P1 *WT* and *Krt6a/Krt6b* null littermates and $5\times10^4$ cells were seeded in a 96-well plate for primary culture. Cells were given fresh media the following day, and labeled 24 hours later with DCFDA dye (2',7' – dichlorofluorescin diacetate abbreviated as "D"; 0-25 μm) and Tert-Butyl Hydrogen Peroxide (abbreviated as "T"; 0-250 μm) to induce oxidative stress. ROS were measured using the Abcam DCFDA cellular ROS detection assay kit. (B) Keratinocytes were cultured from *WT* and *Krt16* null littermates and cultured and analyzed as described in (A). (C) Spontaneously immortalized keratinocytes from P1 *WT* and *Krt16* null mice were seeded at a density of $2.5\times10^4$ cells per well and assayed as described in (A). n= 3 independent experiments. Student's t test; significance at $p<0.05$.

**Supplemental Figure 3** Representative movies of fluorescently-labeled mitochondria in primary cultures of keratinocytes (related to Figure 4). *WT* (Movie 1) and *Krt6a/Krt6b* null (Movie 2) newborn mouse littermates; *WT* (Movie 3) and *Krt16* null (Movie 4) newborn mouse littermates. The quantitative data reported in Figure 4 was obtain from analyzing such movies.

- **Supplemental Video 1.** Representative movie of keratinocytes isolated from the *WT* littermates of *Krt6a/Krt6b* null mice. See main text for details.
- **Supplemental Video 2.** Representative movie of keratinocytes isolated from *Krt6a/Krt6b* null mice. See main text for details.
- **Supplemental Video 3.** Representative movie of keratinocytes isolated from the *WT* littermates of *Krt16* null mice. See main text for details.
- **Supplemental Video 4.** Representative movie of keratinocytes isolated from *Krt16* null mice. See main text for details.

**Supplemental Figure 4.** Complement to live imaging of mitochondrial dynamics (related to Figure 4). Data corresponds to the average mitochondrial speed as a function of mitochondrial profile size for each of three independent experiments involving matched null and WT littermates for each of the *Krt6a/Krt6b* and *Krt16* null lines.

**Supplemental Table 1.** Forward and reverse oligonucleotide primers used for RT-qPCR assays.

| Molecule | Forward (5' – 3') | Reverse (5' – 3') |
|---|---|---|
| *CoX4* | ATTGGCAAGAGAGCCATTTCTAC | CACGCCGATCAGCGTAAGT |
| *Pyruvate Dehydrogenase* | AGGAGGGAATTGAATGTGAGGT | ACTGGCTTCTATGGCTTCGAT |
| *Succinate Dehydrogenase* | GCTGCGTTCTTGCTGAGACA | ATCTCCTCCTTAGCTGTGGTT |
| *Tim23* | GAAGGTGGCGGAAGAAGTAGC | GGGGGTTCATACCAGTCAGC |
| *COR2* | AAAGTTGCCCCGAAGGTTAAA | GAGCATAGTTTTCCAGAGAAGCA |
| *Drp1* | CCAGCTGCCTCAGATCGTCG | CCGGGTGACCACACCAGTTC |
| *Mfn1* | CGTCGAGAACCTCCATGGGC | TCGTCCACGTCAGCCTCTCA |
| *Mfn2* | CTGGGCCCCAAGAATAGCCG | GCGATGAGTCTCCAGCCCAC |
| *Opa1* | AGGGTCCGGCGAGATGATGA | CACCATCCTCTGCAGCCCAG |

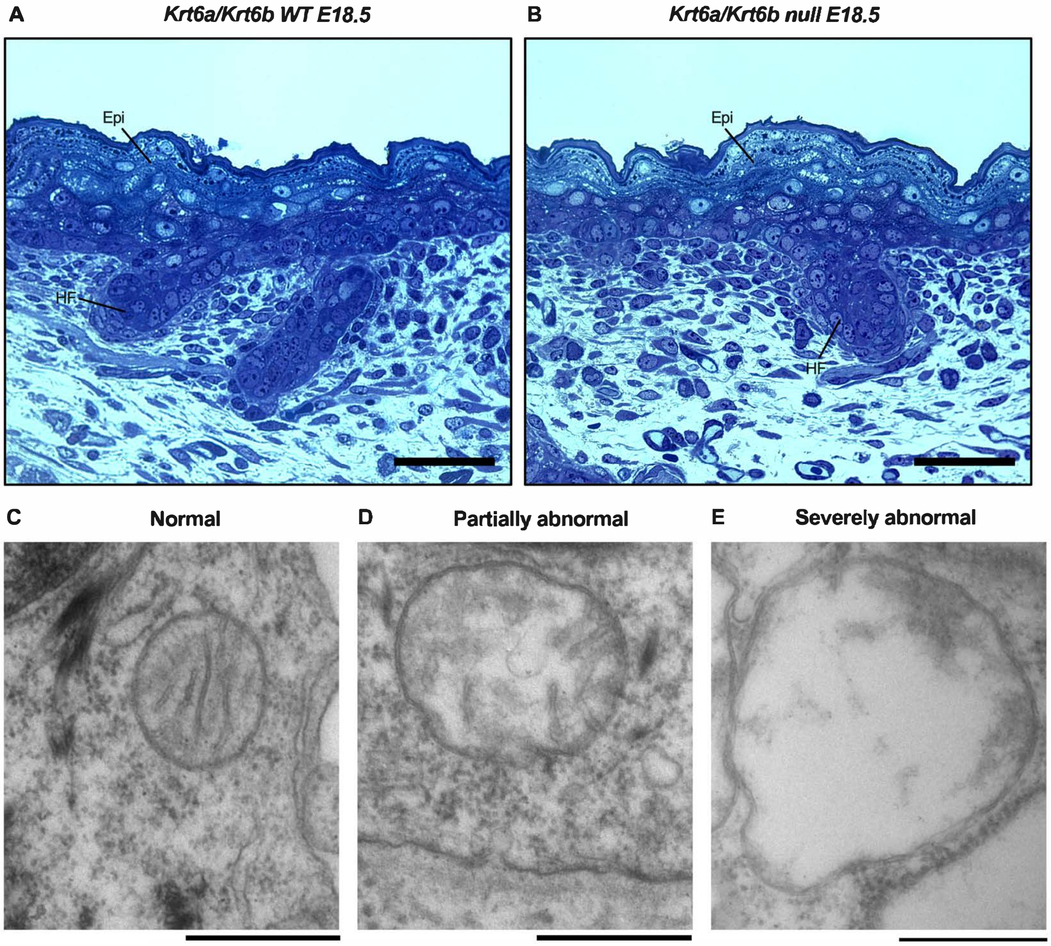


Steen et al., Supplemental Figure 1

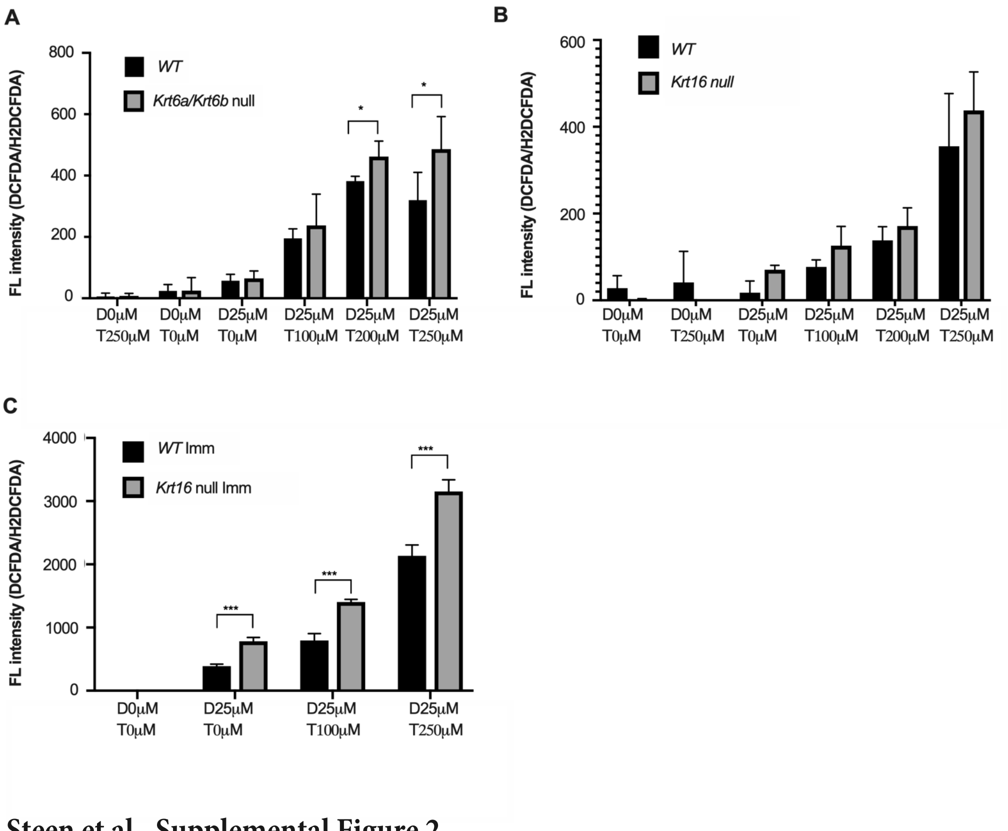


**Steen et al., Supplemental Figure 2**

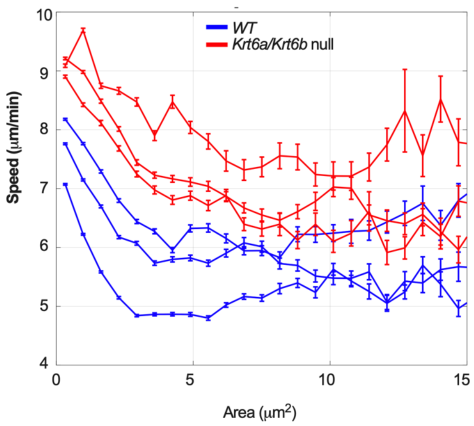


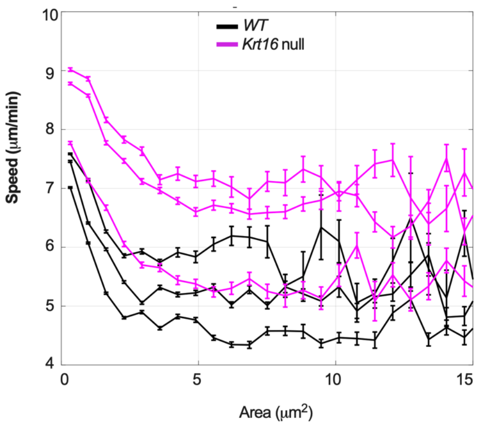


**Steen et al., Supplemental Figure 3**
